# 4D imaging of soft matter in liquid water

**DOI:** 10.1101/2021.01.21.427613

**Authors:** Gabriele Marchello, Cesare De Pace, Silvia Acosta-Gutierrez, Ciro Lopez-Vazquez, Neil Wilkinson, Francesco L. Gervasio, Lorena Ruiz-Perez, Giuseppe Battaglia

**Author notes:** **Corresponding authors:** Dr Lorena Ruiz-Perez and Prof Giuseppe Battaglia,Christopher Ingold Building, University College London, 20 Gordon Street, WC1H 0AJ, London, United Kingdom, and. These authors have contributed equally (Gabriele Marchello, Cesare De Pace).

## Abstract

Water is a critical component for both function and structure of soft matter and it is what bestows the adjective soft. Imaging samples in liquid state is thus paramount to gathering structural and dynamical information of any soft materials. Herein we propose the use of liquid phase electron microscopy to expand ultrastructural analysis into dynamical investigations. We imaged two soft matter examples: a polymer micelle and a protein in liquid phase using transmission electron microscopy and demonstrate that the inherent Brownian motion associated with the liquid state can be exploited to gather three-dimensional information of the materials in their natural state. We call such an approach brownian tomography (BT). We combine BT with single particle analysis (Brownian particle analysis BPA) to image protein structures with a spatial resolution close that achievable using cryogenic TEM. We show that BPA allows sub-nanometer resolution of soft materials and enables to gather information on conformational changes, hydration dynamics, and the effect of thermal fluctuations.

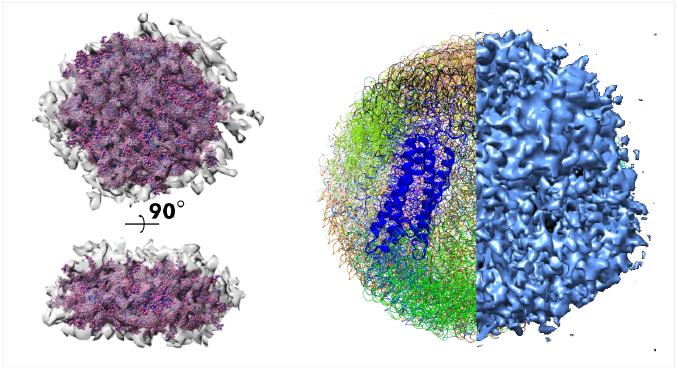

## Introduction

Electron microscopy (EM) is one of the most powerful techniques for structural determination at the nanoscale, with the ability to image matter down to the atomic level. The short wavelength associated with electrons is pivotal for achieving such a high resolution. However, electrons interact strongly with matter making imaging possible only under ultrahigh vacuum. Such a necessity limits EM imaging to solid-state or non-volatile samples. Samples in most common liquids or containing liquids, especially water, thus require special preparation techniques involving either controlled drying or cryogenic treatments with consequent artefacts. Such alterations become particularly critical for biological samples and soft materials whose meso- and nano-scale structures comprise water as a building block. Some of these limitations can be overcome using fast vitrification processes to solidify liquid samples [1], however vitrified water is not liquid water and its structure and hydrogen bond network is very different [2, 3]. Yet, for the last three decades, cryogenic electron microscopy (cryo-EM) has radically changed structural biology [4, 5, 6, 7, 8]. The so called single particle analysis (SPA)[9, 10] in particular enables the atomic reconstruction of almost any type of proteins [11] with tremendous consequences in drug discovery [12].Cryogenic and dry TEM provide snapshots of three-dimensional (3D) structural information that can be correlated with function. The fast combination of different snapshots is however the key to achieve information on structure dynamics. The new frontier of material science, and biology lies in the accomplishment of structure dynamics by including the fourth dimension, time. [13].

The recent development of electron-transparent materials have paved the way to liquid-phase electron microscopy (LP EM) [14]. LP EM imaging of samples in liquid have been reported for electrochemical reactions [15, 16], nanocrystal growth [17, 18], whole cells[19, 20], and tomography reconstruction of particles in liquid state [21]. Soft materials such as micelles and vesicles form in water, or any other selective solvent, as a result of counteracting interactions including hydrophobic effects, steric hydrophilic interactions, hydrogen bonds and electrostatic interactions [22]. Such relatively weak forces give rise to highly dynamical assemblies that exist in solution where the solvent plays a critical role in the integrity of such structural assemblies. LP EM is thus becoming the ideal tool to study the evolution [23] and formation [24, 25] of vesicles and micelles in solution. In LP EM, the 3D reconstruction of particles in liquid has been mainly dominated by the analysis of inorganic particles [21, 26]. While SPA methods have been used to obtain the 3D structure of proteins (by entrapping them through substrate functionalization) [27], protein dynamics has been characterized only indirectly with these approaches.

In our investigation, we exploited LP EM to image the dynamics of particles undergoing Brownian motion, using their natural rotation to access all the particle views in order to reconstruct their 3D structure using tomographic techniques. We propose two distinct approaches to assess the performance of the employed tomographic techniques for reconstructing 3D soft structures from LP EM imaging.

The first approach involves the reconstruction of heterogeneous particles while in the second approach we reconstruct homogeneous ones. For heterogenous populations, we used the Brownian tomography (BT) method based on the analysis of a single unit that is imaged long enough to access all the views of its whole structure. For homogeneous populations such as globular protein dispersions we present the Brownian particle analysis (BPA) where BT is combined with SPA methods to allow for 3D reconstruction with sub-nanometer resolution as a function of time.

## Results and discussion

### Liquid phase electron microscopy of soft materials

In order to assess the ability of LP EM to image proteins and synthetic soft materials in liquid phase using direct detection and fast acquisition tools we chose the protein ferritin as model. Ferritin is a globular protein complex comprising 24 chains arranged into a rigid and very stable tetracosameric structure with octahedral symmetry, forming a hollow spherical shell [28]. Ferritin is a nanoscopic cage containing multiple Iron ions with an external and internal diameter of 12 and 8nm respectively, already imaged via LP EM [29]. We imaged ferritin dispersed in both deionised water, and phosphate buffer solution (PBS) using a complementary metal-oxide-semiconductor (CMOS)-based direct electron detection camera (Gatan K2 IS) which allows low-dose imaging with high detective quantum efficiency (DQE) [30]. The employed direct detection device (DDD) allows two Electron Detection methods; Linear, where all the detected electron charge in the sensor is integrated into the resultant image; and the counted Mode, where the resultant electron interaction is processed to a single count per electron event. With the very fast speed of the DDD, images are collected in Dose Fractionation mode, where multiple images can be collected as a movie in place of a single exposure, thereby giving access to the time dimension. However, the imaging processes associated to LP EM generates particularly noisy images due to the liquid media[32]. We assessed that contrast can be improved by imaging the ferritin dispersion with a negative defocus and sample height values as shown in **Fig. S3** where the proteins appeared as single bright dots encircled by a dark ring. All together these modalities help to image ferritin in liquid phase (**Fig. 1a**) reaching contrast similar to that afforded by cryogenic fixation **Fig. 1b** under the same imaging conditions. We firstly observed a critical difference in sample resistance to the electron beam when using either DI water or PBS as solvent for the ferritin dispersions. In **Fig. 1c** we show a sequence of ferritin dispersed in DI water at 2.5mg/ml imaged over 4 seconds at a dose of 30 electrons Å^−2^*s*^−1^. The resulting images show a quick degradation and consequent morphological changes of ferritin which upon close inspection seems to form large aggregates with structural similarities to amyloid fibrils. We analyzed the particle size in each of the forty frames in an attempt to unveil morphological changes as a function of time. The relative size distribution of ferritin seemed to change from an average of 12 nm (the ferritin outer diameter) to larger and more dispersed structures (see **Fig. 1d**) at the end of the four seconds of acquisition time. Inversely, when ferritin was dispersed in PBS at the same concentration as that for DI water, we were able to image ferritin at a higher electron dose (140 electrons per Å^2^ per seconds) with no indication of sample damage, as shown in **Fig. 1e**. We deliberately chose to image proteins that were located close to the SiN window and hence did not seem to physically translate much during imaging with the aim of maximising their cumulative electron dose. Not only did we observe no changes in the morphology of ferritin, but its average particle diameter stayed constant at 12nm over the four seconds of acquisition. (see **Fig. 1f**). The total electrical charge in the specimen illuminated area can be estimated [33, 34] as *Q* ∝ *I*_*S*_*ϵϵ*_0_ *σ*^−1^ where *I*_*S*_ is the specimen beam intensity, *σ* and *ϵϵ*_0_ are the medium conductivity and relative permittivity respectively. Deionised water, when used as the medium for dispersing ferritin, has a conductivity at room temperature of *σ*_*DW*_ = 5×10^−6^*Sm*^−1^, while the PBS used in our experiments is a water solution of potassium dihydrogen phosphate, sodium dihydrogen phosphate and sodium chloride and accounts for a conductivity of *σ*_*PBS*_ = 2*Sm*^−1^. Assuming that *I*_*S*_ is the same for both media, and with water relative permittivity *ϵϵ*_0_ = 80, the specimen charge in deionised water will be 4 × 10^5^ higher than in PBS. Such a difference in specimen charge will be even more enhanced when compared to water in its vitrified state, as in Cryo Electron Microscopy. In the latter case, depending on the relative permittivity and *I*_*S*_, and due to the low conductivity of vitrified water, we would expect ferritin to have a much higher charge, between one to three orders of magnitude in vitrified than in liquid water. In the case of proteins, and in particular ferritin, significant implications arise as a consequence of the reduction of charge associated with the liquid media, allowing to effectively increase the lethal beam illumination/dose threshold for protein samples when they are imaged via LP EM.

**Figure 1:**
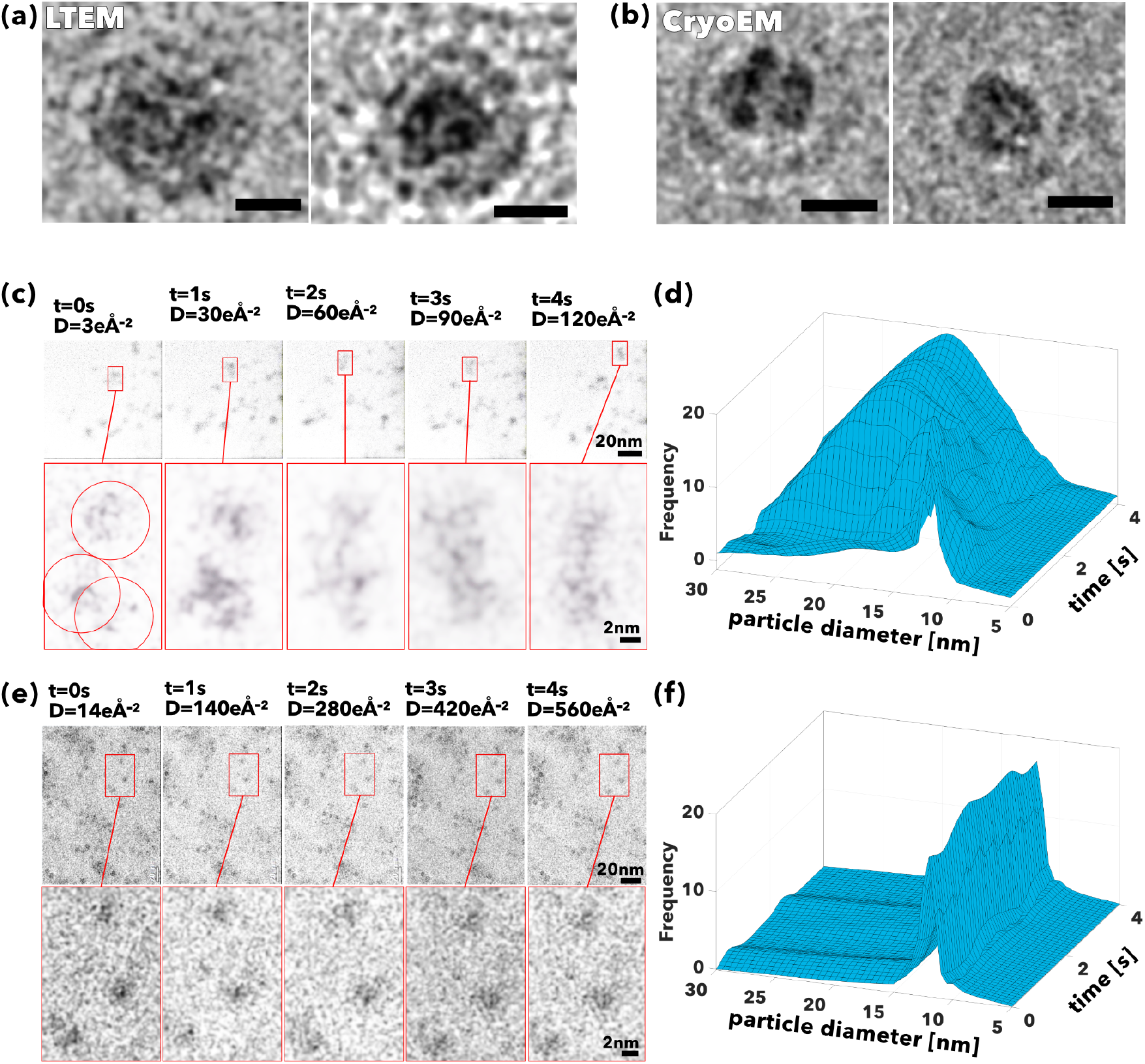
Electron dose and contrast in LP EM. TEM micrographs of ferritin proteins imaged in liquid **(a)**, and vitrified **(b)** water TEM methods. LP EM micrographs and particle size distribution as a function of time of ferritin proteins dispersed in deionised water **(c)** and **(d)** and phosphate buffer solution **(e)** and **(f)** imaged over 4 seconds at 10fps acquisition **(a)**.

### Brownian tomography

Having assessed the optimal imaging conditions in terms of electron dose, we proceed to assess the ability of LP EM to image dynamical changes. Here we chose to work with block copolymer micelles formed by poly(2-(methacryloyloxy)ethyl phosphorylcholine),Äìpoly(2-(diisopropylamino)ethyl methacrylate)(PMPC,ÄìPDPA) amphiphilic copolymers in water. We recently reported[35] that the PMPC-PDPA copolymer can assemble into disk-like micelles with a thickness of about 15nm and a radius from up to 30nm. These metastable structures emerge from the bottom-up assembly of membrane-forming copolymers when the system is quenched before the critical number of aggregates to form vesicles is reached [35]. We previously imaged the micelles in both dry and cryogenic conditions[35] observing that in both cases, the micelles tend to bias their orientations showing preferentially in most cases the widest side, hence hindering their full characterisation. In liquid mode, as shown in video1 in the SI, the micelles are free to rotate and move with no preferential orientation hence displaying all their views. As shown in **Fig.2**, we can image a single micelle in time allowing the Brownian translation and rotation to randomise its orientation. In **Fig.2a**, we show four different snapshots of the micelle orientation over time provided by LTEM .The difference observed in the various views acquired sequentially does indeed verify the anisotropy in the geometry of the disk like-micelle. Each view is completely random as the result of Brownian rotation, as showed by the time-coloured palette in **Fig.2b** revealing the stochastic evolution of each view as a function of the imaging time. Unfortunately, our temporal resolution is 100ms, four orders of magnitude longer than the rotational relaxation time of the investigated structure. The differences between the afforded temporal resolution and rotational relaxation times hinders an effective tracking of the angular dynamics. Yet, with the Brownian Motion and the fast image acquisition rate, we can capture a large number of views and use them sequentially to reconstruct the full 3D structure. This latter is shown in **Fig.2c** as an axonometric projection of the disk like micelle alongside the different orientations monitored in red and three of its views in **Fig.2d** effectively creating what we call a “Brownian tomography” of the micelle.

**Figure 2:**
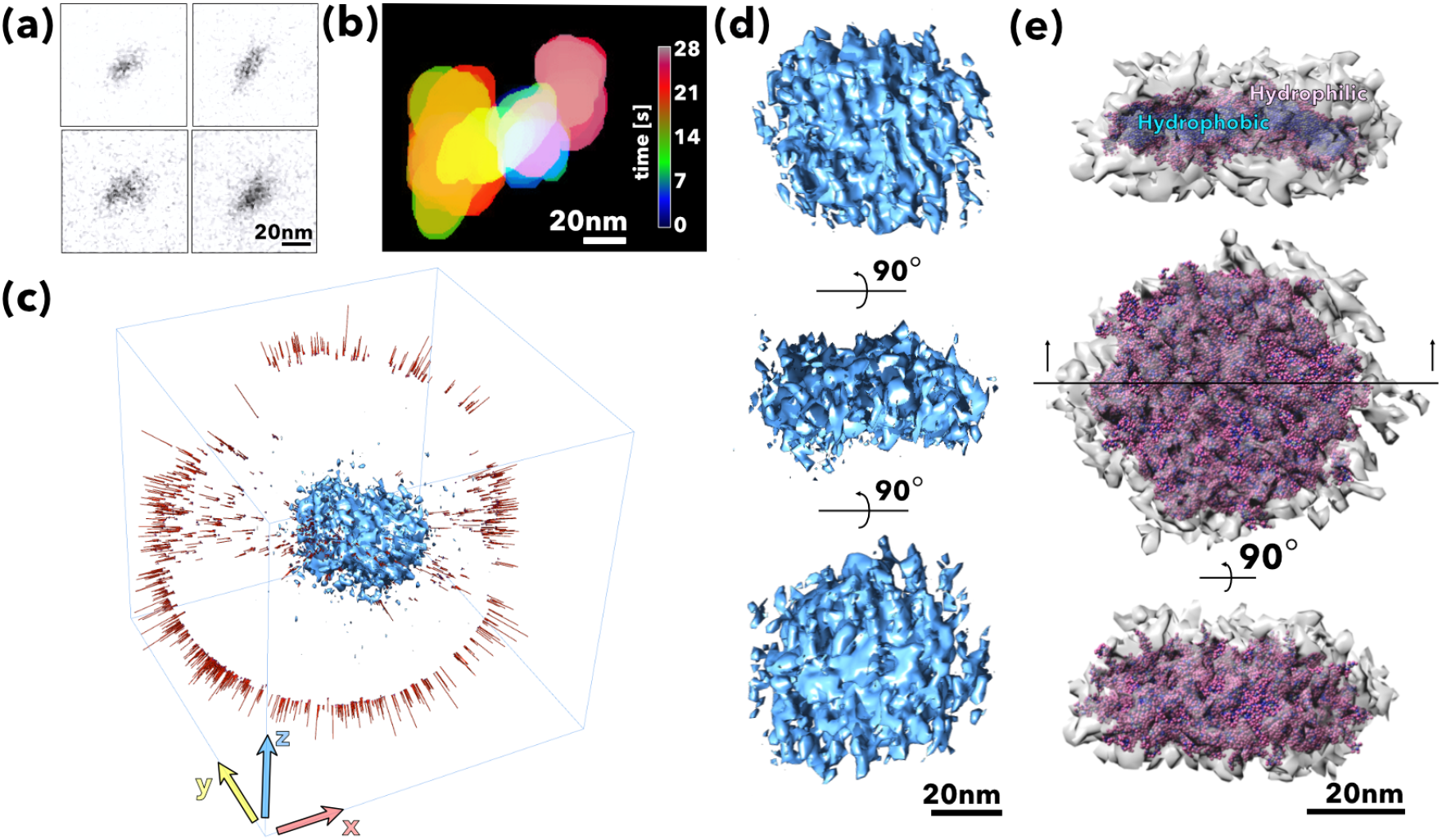
Brownian tomography of a block polymer disk-like micelle. Snapshots of a single PMPC-PDPA disk-like micelle rotating in water imaged in liquid phase transmission electron microscopy **(a)**. Time-coloured palette sequences of a PMPC-PDPA disk-like micelle imaged sequentially **(b)**. Each colour represent a different time point. Axonometric projection of a PMPC-PDPA disk-like micelle shown alongside the different orientations collected as a function of time **(c)**. Three dimensional rendering of a PMPC-PDPA disk-like micelle shown in three different views **(d)**. Comparison between the coarse-grain model and the Brownian tomography rendering shown in two orientations and as cross section showing the hydrophobic core surrounded by the hydrophilic brush **(e)** .

It is evident that we can visualize the entire micelle structure with a resolution of circa 1nm almost allowing the visualization of its polymer building blocks. its polymer building blocks. Such a level of resolution allows us to extract further structural information from the micelle by combining our experimental data with computational modelling. As shown in **Fig.S2a**, we can “coarsen” the single PMPC-PDPA chain in different units or beads that comprise the critical chemical elements, allowing us to reduce each polymer from 2536 atoms to 316 beads. The use of the coarse model allows us to simulate the full disk (1million atoms) and capture the full dynamics of the system including its self-assembly behavior. The resulting structure can then be compared with our Brownian tomography by fitting the number of polymers as free parameter as shown in **Fig.S2b**. The result shows a very good agreement between the experimental Brownian tomography and the coarse-grain simulation capturing the different structural elements (see **Fig.S2c**) of the micelles. In addition, the proposed methodology allows us to extend our morphological analysis down to the single chain level. We assume now that the end-to-end distance for each polymer segment is *d* ≃ *a* ∗ *N* ^*ν*^ where a is the monomer length, *N* is the polymerisation degree, and *ν* ∈ [0, 1] is the conformation scaling factor. From polymer physics[36], we know that an unperturbed polymer chain scales with *ν* = ⅗, when the chain is fully stretched in ideal conditions *ν* = 1, while when the chain is compressed and hence coiled up *ν* ≤ ⅗. The coarse-grain simulations show that PMPC chains are all fully stretched in water with an average *ν* ≃ 1 while the PDPA hydrophobic chains show an end-to-end distance that is dependent on their physical location within the micelle. As shown in **Fig.S2d**) and as expected by our disk formation theoretical model[35], the chains in the middle of the structure are unperturbed or slighlty streched[37], while the chains on the disk edge are collapsed and coiled up.

### Brownian particle analysis

We have shown that LP EM can be used to capture sufficient particle views to create 3D rendering of nanoscopic objects undergoing Brownian motion. We further extend the capabilities of LP EM by its application for evaluating protein structure. Following the same reasoning as introduced above for the disk-like micelle, proteins are free in liquid media and hence undergo Brownian rotation and translation with each protein changing stochastically its orientation over time. Thus, imaging tens of proteins over few seconds allows us to capture thousands of protein views. We exploit this feature using the SPA algorithm but proposing here the Brownian particle analysis (BPA) approach. While SPA requires a great number of particles in solution to be immobilized on a grid in order to preserve their structures under vacuum[38], BPA has the ability to process videos that may not contain thousands of particles, but instead monitor individual particle views across many frames, rather than across a single frame. In this fashion, the orientations needed for a successful 3D reconstruction are produced by recording the motion of the particles in solution. Motion in this context refers to Brownian motion in confinement due to the physical characteristics of the liquid cell. It is not the scope of the present study to distinguish between the motion experienced by the particles when they are located in interfacial areas close to the observation windows or in the bulk. We exploit the motion of the particles for the sole purpose of visualizing the particles views.

Time now plays a pivotal role in the reconstruction process and a few seconds of videos can replace hours of sample preparation and complex data collection necessary for cryo-EM. The video 2 in the supporting information was processed employing the BPA reconstruction technique and it displayed a few hundreds of ferritin particles in each frame. Video 2 was recorded for 10s at 10fps, generating a hundred frames. Over 5000 views of ferritin were extracted from the first fifty recorded frames (*i*.*e*. over the 5s video recording) and used to reconstruct the density maps. The core of the reconstruction algorithm was performed with Scipion, freeware designed to perform SPA 3D reconstruction, that was extended and adapted to our needs[39]. Firstly, the stack of two-dimensional (2D) micrographs was pre-processed to reduce the noise using a median filter. The ferritin views were identified, extracted across all the frames, and collected together to generate a 3D density map of the ferritin protein. As detailed in **Fig. S4** a single frame contains circa 100 proteins per field of view that can be tracked over time thus allowing the imaging of sequential views in consecutive frames. The BPA method can be considered a reference-free technique, because it does not use any *a priori* information about the structures of the imaged particles. A coarse 3D density map was initially reconstructed, and the map was subsequently refined by applying the projection matching algorithm to produce an unbiased, high-fidelity density map as shown as front view and equatorial cross section in **Fig. 3a**. The reconstructed density map was further validated by using the ResMap algorithm to measure the local resolution [40] providing a mean and median resolutions of 4.7Å and 4.8Å respectively (**Fig. 3b**).

**Figure 3:**
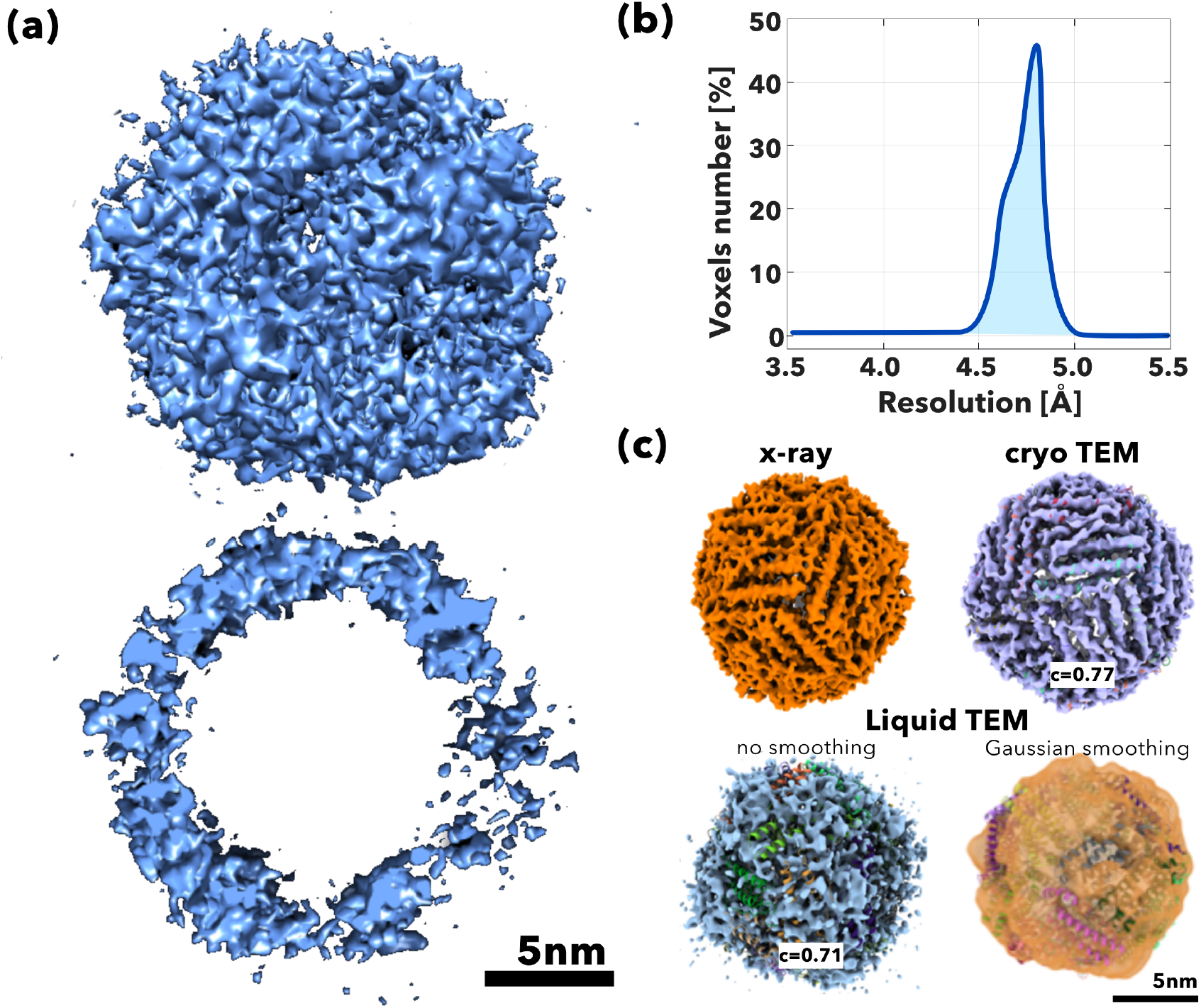
Brownian particle analysis. Ferritin density map calculated using the Brownian particle analysis method, show as front and equatorial view **(a)**. Corresponding local resolution of the ferritin density map performed through ResMap **(b)**. Comparison between x-ray, cryo-EM and Liquid TEM shown smoothed and Gaussian-smoothed density maps **(c)**.

**Fig. 3c** shows the electron density (ED) map generated from x-ray structure (PDBid 6MSX), the cryo-EM (EMD-2788) map reported by Russo *et al*.[41], and the LTEM ED maps from **Fig. 3a** displayed with no smoothing and Gaussian smoothing applied. All ED maps are shown overlaid with the ferritin atomistic structure from x-ray with the different alpha helixes represented in green, yellow and red. In order to quantitatively evaluate the differences in the ferritin ED maps obtained by the various methods, we employed a rigid-body fitting of the PDB 6MSX with the ED maps using ChimeraX [42]. The ED map and supramolecular complex were superimposed to maximize the number of atoms inside the map followed by a rigid-body local optimization using a map-in-map fitting procedure maximizing the correlation value, *c*, between the map and the reference[43]. We measured for the cryo-EM (EMD-2788) ED map a correlation of *c* = 0.77, while for the LP EM ED map generated over 5s video recording a correlation of *c* = 0.71 with the x-ray reference map.

One of the major advantages of LP EM is that the ferritin ED map is the result of sequential views and thus we can break it down to assess the evolution over time. We thus segmented the same video used for the reconstruction into five consecutive, not-overlapping intervals, containing ten frames each. These five one second-long video segments were processed, and the associated 3D density maps were reconstructed. Each video segment contained about 1000 different protein views. The apparent low number of particles could lead to low-quality density maps, however both coarse and refined maps (**Fig. 4a**) yield useful information about the evolution of the particles in time. Although the particles being the same type should lead to five identical 3D density maps, different configurations of the maps surface and shell outer contour were obtained. The ResMap algorithm measured resolution mean values of 5.42Å, 5.39Å, 5.39Å, 5.37Å, 5.37Å; and median values of 5.5Å, 5.4 Å, 5.4 Å, 5.4 Å, 5.4 Å, respectively (**Fig. 4**b). The differences amongst the local resolution distributions, particularly for the first video segment and the long-exposure density map accounted for the difference in the number of particles involved in the reconstructions. Similarly, the comparison of each map with the x-ray reference map shows similar values of correlation with the second ED map corresponding to the time interval between 1 and 2 seconds producing an even higher correlation value of *c* = 0.75 as shown in **Fig. 4c**. In the same graph we plotted the cumulative electron dose as a function of time, maintaining beam illumination constant during acquisition, and indeed it shows that the cumulative dose increases up to 700 electrons per Å^2^ [44]. Whilst a longer acquisition time for the movie can generate more particle views, the increase in exposure may lead to beam damage, which needs to be considered. The analysis of the resolution obtained from the five time-varying maps leads to some interesting considerations. The number of missing patches on the density maps seemed to increase over time, while the resolution of the density maps remained constant. In order to have a reference for the quality of the images in terms of SNR, and reconstructions for different magnitudes of electron doses we repeated the same experiments with a cumulative electron dose of less than 10 electrons per Å^2^. The resulting ED maps are shown as no binned, and binned over 2, 5 and 10 frames in **Fig.S5a**. Whilst frame binning compromises on time resolution, as the interval needed to capture high number of views becomes longer, it improves the quality of the reconstruction compared to not-binned map by an improved signal noise ratio. We compared the low dose binned maps with the reference, and as shown in (**Fig.S5b**) the resulting ED maps appear to have missing surface patches. The 3D map obtained after binning twice the frames in the low dose video showed a correlation value of c=0.66. Increasing the binning size seemed to decrease the map correlation values (**Fig.S5c**). Most importantly, within the limited resolution, the overall structure observed at low dose is similar to that obtained from high electron dose, thus confirming again the improved beam resistance of ferritin when dispersed in liquid media. With this in mind, the density map generated from cryo-EM has sufficient resolution to distinguish the *α*-helices represented in green, yellow and red. The generated density map from LP EM is coarser than the map generated from cryo-EM and even though our ResMap calculations suggest spatial resolution below the *α*-helices dimensional features, we cannot distinguish these features. It is worth mentioning that ferritin remained intractable to structure determination by cryo-EM for a long time due to the fact that the contrast afforded by the individual imaged particles was insufficient for alpha-helices to be resolved. *α*-Helices resolution is required for aligning the images to each other and thus superimpose the imaged structures [44]. This problem was first solved by using a gold support and selecting the images of proteins with reduced motion in the gold grids [45]. Herein we capture the protein structure at room temperature in liquid water with rotational correlation time *τ*_*R*_ ∼ 1*μs* but also with internal structure vibrations with correlation times in the order of ns. Hence, our reconstructions, independently of time intervals used, are time-averaged and represent the volume occupied by all the protein conformations within the time scale used for imaging (∼ 100*ms*). In order to elucidate the possible role of the protein dynamics in the observed frames and reconstructions, we performed long molecular dynamics simulations of ferritin in solution (150mM NaCl). The ferritin supramolecular complex is very stable and retains its tetracosameric structure along simulation time (250ns). In **Fig.5a** we report the pairwise root mean square deviation (RMSD) matrix for the ferritin complex. There are very few differences in conformations (as measured by RMSD) across the trajectory, meaning the complex samples similar conformations. In **Fig. S7** we report the single chain RMSD distributions for the 24 ferritin chains labelled from A to X. We observe a RMSD of ∼ 1.5 Å in the simulation which is relatively small for such a large complex, with chains D, M, and U fluctuating more than the rest yet without any major rearrangements. The supramolecular complex is very rigid and lacks significant dynamical changes; if we overlay the MD simulation of all the 24 chains forming ferritin together as in **Fig/ 5b** the result is a blurred structure where it becomes impossible to distinguish the chains secondary arrangements. However, if we overlap the MD overlay with the LP EM ED map, the correspondence is good. This optimal match indeed suggests the key ability of LP EM is to access the conformational spatial flexibility of the protein hence capturing the protein dynamics together with conserved structural features, which for the ferritin is its spherical core shell structure

**Figure 4:**
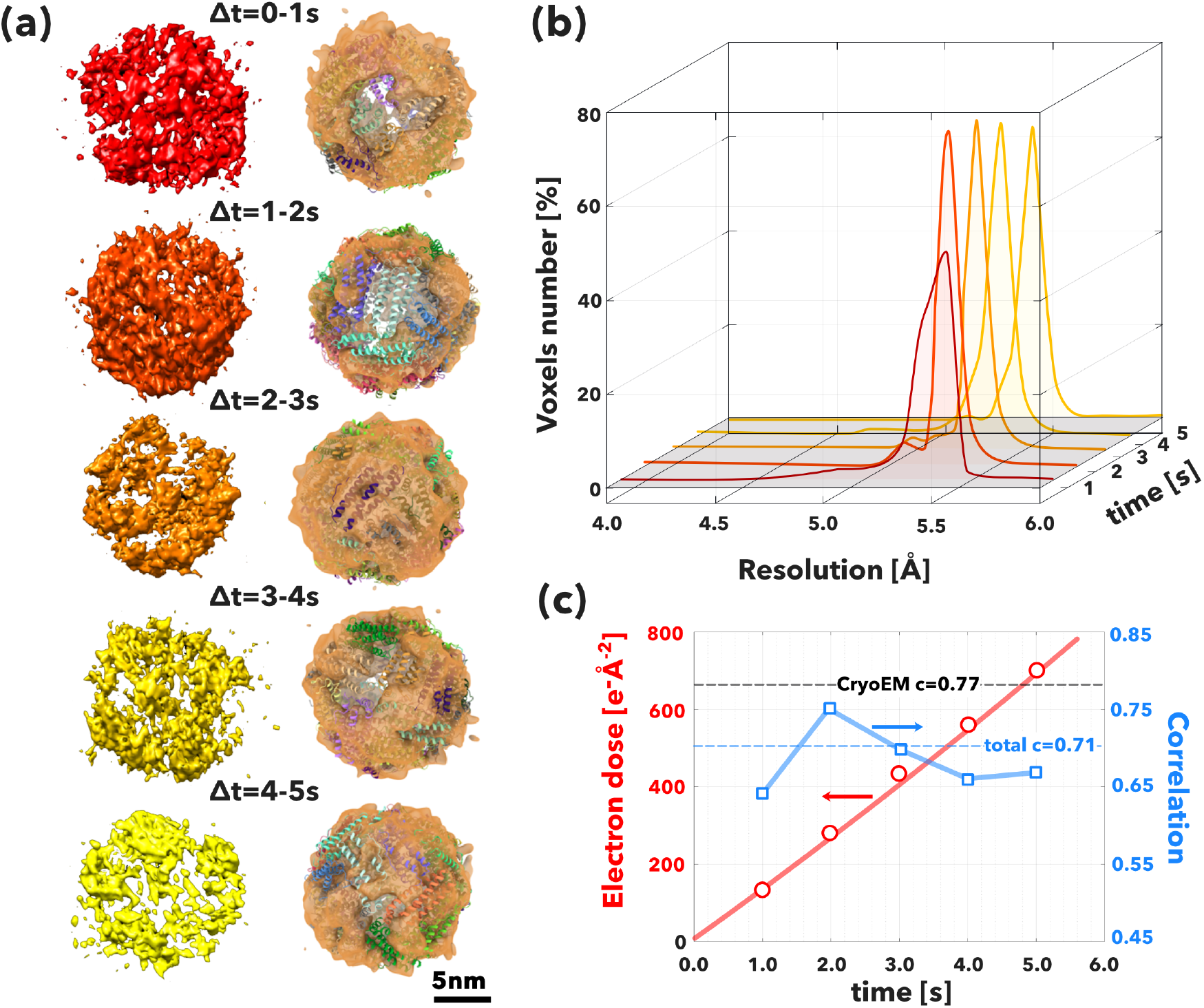
Dynamic reconstruction of ferritin as a function of time. **(a)** Schematic representation showing the temporal evolution of the density map reconstruction process. The algorithm extracted about 1000 particles view from each sub-video, generating five different density maps shown as course and refine the model. **(b)** The histograms generated from the measurements of the local resolution of each density map performed through ResMap for each temporal segment. **(c)** The electron dose the particles were exposed to expressed for each 3D density map and plotted alongside the mean spatial resolution.

**Figure 5:**
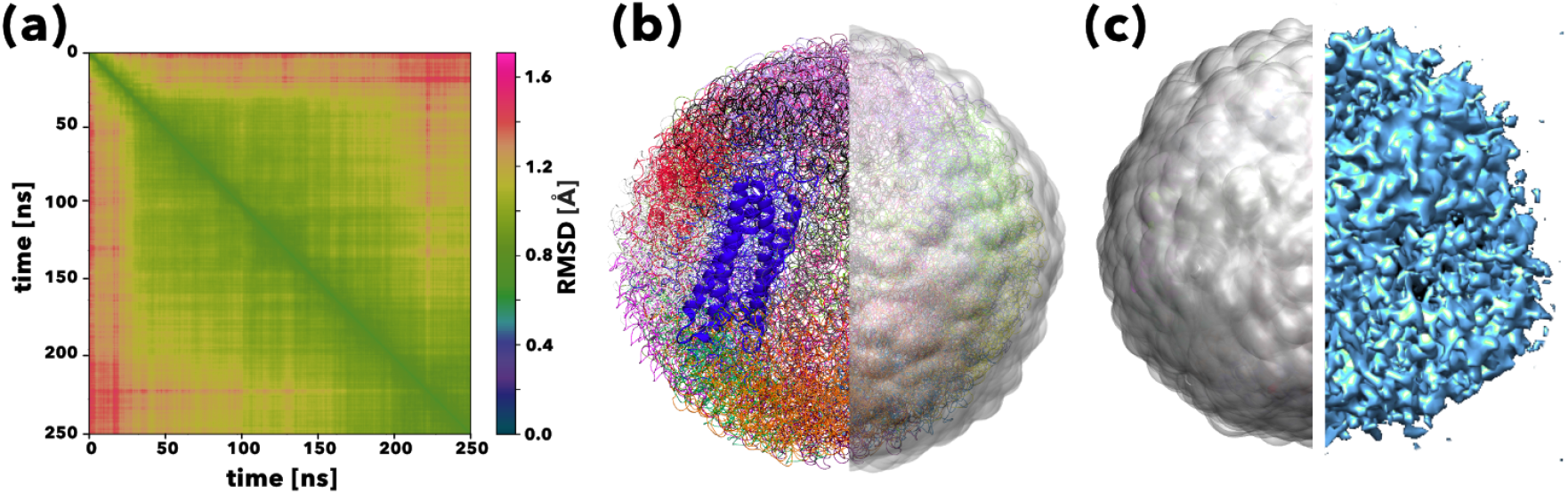
Imaging the protein conformational space. Root mean square deviation pairwise matrix for the ferritin supramolecular complex across the molecular dynamics trajectory **(a)** calculated from Molecular dynamic simulations of horse-spleen ferritin (PDBid 6MSX) light chain depicted as cartoon with one half showing the corresponding overlaid electron density map in gray **(b)**. Overlay of the ferritin conformation analysed by molecular dynamics across 100ns comparison between the LTEM ED map and the ferritin conformational ensemble **(c)**.

## Conclusions

We demonstrate that liquid phase TEM can be used effectively to image soft matter and proteins, in particular capturing their dynamical motion in water. The extra temporal dimension allows us to convert the Brownian motion associated with the particle diffusion into tomography techniques that can screen the full three dimensional view of the specimen. We also showed that Brownian tomography can be combined with single particle analysis techniques to efficiently perform a “Brownian particle analysis” to reconstruct protein structure in liquid water at room temperature with resolution close to benchmarks methodologies. We show that we can reconstruct 3D density maps of protein in liquid solution. The time factor becomes a fundamental element in the reconstruction process, reducing the acquisition time required to collect enough particles from hours in cryo TEM to seconds. In time, with the evolution of detector technologies, the resolution will increase, and avoiding the vitrification process will allow interesting comparisons. Most importantly, BPA allows extending atom level structural biology into dynamic structural biology complementing liquid phase NMR and single molecule spectroscopy with a set of unique strengths. Such unique strengths are thus able to shed light on real time conformational changes in solution, hydration dynamics, or merely the evaluation of the effects of thermal fluctuations. We believe that our findings pave the way to a completely new frontier in soft matter and structural biology thus expanding functional studies to dynamics where we can sample soft particles and proteins within a three-dimensional plus time *i*.*e*. 4D space. Such a space will allow us to understand conformational changes, allosteric interaction, folding-unfolding mechanisms as well as to bring the so far neglected role of water in protein structure.

## Acknowledgements

We acknowledge the EPSRC (grant EP/N010906/1) and Jeol UK and DENSsolutions for sponsoring part of this work, LRP salary as well as GM and CDP scholarship. GB acknowledges the EPSRC for funding part of his salary via an Established Career Fellowship (EP/N026322/1) and the ERC (CheSSTag 769798) via a consolidator award F.L.G. acknowledges funding from EPSCR (Grants EP/P022138/1, EP/P011306/1) and European Commission H2020 Human Brain Project CDP 6. HEC-BioSim (EP/R029407/1), PRACE (BSC, Project BCV-2019-3-0010 and CSCS Project S847), CSCS (Project 86), and the Leibniz Supercomputing Center (SuperMUC, Project pr74su) are acknowledged for their generous allocation of supercomputer time.

## Author contributions

G.B. and LRP ideated the work, supervised the experimental work and imaging analysis. CDP and LRP produced and analyzed all the experimental data. NW advised and supported the imaging acquisition. GM analysed and processed the images as well as rendered the different 3D models. CLV produced and purified the disk like micelles. SAG and FDG designed, performed, and analysed the molecular dynamics simulations. All authors wrote and reviewed the manuscript

## Data availability statement

The data the support the finding of this study are available from the corresponding authors upon reasonable request

## Methods

### Materials and Liquid TEM

Polymers were obtained through ATRP synthesis as previously reported [35]. Equine spleen apoferritin was purchased from Merck (previously Sigma-Aldrich) and diluted with phosphate buffer solution (PBS) at a concentration of 2mg/ml. First, the chips were rinsed in HPLC-graded acetone followed by isopropanol for five minutes each, to eliminate their protective layer made of poly(methyl methacrylate) (PMMA). Then, the chips were plasma cleaned for thirteen min to increase their hydrophilicity. Soon after, the chip with the spacer was fitted at the bottom of the liquid cell and 1.5Œ°L of ferritin solution was deposited onto the window of the chip. It was important not to let the ferritin solution dry at this stage as the film thickness of dried ferritin solution cannot be easily controlled and impedes good imaging conditions. The chip without spacer was then positioned on top of the liquid layer closing the liquid cell. The liquid holder was sealed, and 300Œ°L of the ferritin solution at 2mg/mL was flushed in at 20Œ°L/min with a peristaltic pump. A volume of 300Œ°L of solution ensured that the holder tubing system and cell were filled with ferritin solution. We waited five minutes after collecting ferritin solution from the outlet tube (Fig. S1) in order to guarantee that convection effects from the flowing process were not affecting the Brownian movement of ferritin in solution. The peristaltic pump was then stopped and the inlet and outlet tubes were sealed to ensure a closed liquid circuit. The liquid holder was then introduced into the microscope. Imaging was performed with the holder in static conditions i.e. the peristaltic pump was stopped during imaging. Fluid dynamics simulations were performed on the liquid holder and are reported in the section addressing Fluid dynamics simulation.

LP EM imaging procedure. The experiments were performed using a JEOL JEM-2200FS TEM microscope equipped with a field emission gun (FEG) at 200 kV, and an in-column Omega filter. The camera used was the direct detection device (DDD) in-situ K2-IS camera from Gatan. The ultra-high sensitivity of the K2 allows low-dose imaging modes limiting considerably electron dose damage and facilitating high spatial (3838×3710 pixels) and temporal (up to 1600 fps in in-situ mode) resolution. The microscope was used in transmission electron microscopy mode. In order to limit the beam dose on the specimen, images were collected at the minimum spot size (5) with a small condenser lens aperture (CLA 3). The K2 can be used in two recording modes, linear and counted. The former is fast, but it generates the output image by capturing the integrated signal from each pixel, I n a similar way to a standard charge coupled device (CCD). Counted mode processes the image and assigns one count to the pixel or sub pixel of the landing site of that electron, thereby illuminating background noise. For our investigations dose fractionation videos were recorded in linear mode. Further details on the quality of the videos are provided in section Video analysis. The electron flux used to record the videos was 14 e-Å^−2^, that is 140 e-Å^−2^. Low electron dose videos were also recorded at circa 0.9 e-Å^−2^ under identical imaging conditions as the high dose 5s video. Underfocus of circa 10*μm* was needed to improve the contrast of ferritin as phase plates were not present in our imaging system. Every image was recorded in the format of 4-byte grayscale and required the full size of the detector. However, the images were binned by a factor 2 on both dimensions, resizing images down to 1919×1855 pixels. This process not only increased the signal-to-noise ratio (SNR) of each processed frame, but also prevented the memory of the processing machine to saturate due to the high volume of data

### Electron density map comparison

Briefly, we define the ED map as the vector **u** and the reference map, calculated from the x-ray structure, as the vector **r**, the correlation, *c*, between the two maps is thus defined as the ration between the two vectors inner and norm products,

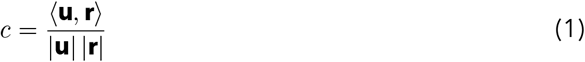

### Molecular Dynamics: Ferritin supramolecular complex

We simulate ferritin starting from the biological assembly of a 1.43 √Ö resolution x-ray crystal structure (PDB-id: 6MSX) containing Cd2+. Only the positions of the protein coordinated Cd2+ atoms were retained and simulated as Fe2+. The protein complex presented no mutations or structural gaps; therefore, it was protonated at pH 7.4 and solvated (150mM NaCl) using a TIP3P water model and Charmm36m(4) protein and ions force field. The final simulated system consisted of 336806 atoms, resulting in a truncated octahedron simulation cell of 16.14nmx15.21nmx13.17nm (5.37, -5.37,7.6). All simulations were performed with Gromacs v2019.2 (5) with a 2fs integration step. We performed 2000 steps of of minimization (steepest descent algorithm), followed by 5000 steps of isotropic NPT at 300K, using the V-rescale thermostat and Parrinello-Rahman barostat. After 50000 steps of NVT the system was fully equilibrated and ready for production that was performed in the NVT ensemble for 370 ns. The last 250ns of the molecular dynamics run were analyzed in order to characterize ferritin dynamics. RMSD pairwise matrix was calculated with MDAnalysis python package [ref 1-5].

### Molecular Dynamics: PDPA72-PMPC25 disk

The PDPA72-PMPC25 all-atom (2536 atoms) structure was drawn using Marvin 19.3.0 (Chemaxon, http://www.chemaxon.com) and the minimum energy conformer was calculated using Marvin Calculator Plugins using a semiempirical force field (MMFF94). A schematic illustration of the coarse graining strategy for PDPA and PMPC in combination with the chemical structure in show in **Fig.S2a**. The coarse-grain strategy for the PDPA72-PMPC25 polymer (316 beads) is based in the recently published model for PDPA (6). All the interaction parameters between groups are identical to the values presented in ref (6), with a reference bond length of ro=0.47nm and a spring constant kb=8000 kJ mol -1 nm-2, based in the MARTINIv2.2 force field beads definitions. A total of 408 polymers (128928 beads) where assemble starting from a sparse bilayer-like conformation (Figure S7) into a compact bilayer by alternating anisotropic and semiisotropic Berendsen barostat (keeping the pressure in the axis normal to the bilayer plane equal to zero). The final frame was manually cut into a disk (r=19nm) and solvated with MARTINIv2.2 non-polarizable water. The final simulation box contained 353 chains of PDPA with a degree of polymerization N=72 linked each linked to a chain of PMPC with a degree polymerization N=25 and 309204 water beads. The solvated system (Figure SX7, 420752 beads) was simulated in the NPT ensemble with an isotropic Parrinello-Rahman barostat for 1 ns followed by a production run of 100ns in the NVT ensemble. All simulation where performed with Gromacs v4.6.7 (Gromacs) software package. The temperature in all production run was T=300 K with a Berendsen thermostat.

## 1 Supporting information

### Liquid cell preparation and assembly

The holder used for imaging ferritin in liquid state was the Ocean liquid holder manufactured by DENSsolution. The ferritin in solution was sand-wiched in a liquid cell formed by two chips made of silicon nitride (SixNy), with 50nm thick rectangular observation windows at the centre measuring 20mmx200mm. Silicon nitride prevents the solution from leaking from the closed cell and allows the passage of some of the electrons through the sample as it is electron transparent. The liquid cell is comprised of a spacer-chip, placed at the bottom and a spacer-less chip placed at the top. 200nm spacer chips were used for the reported investigation. Although the liquid thickness is dictated by the spacer used in the liquid cell, the observation windows of the chips tend to experience some bulging, mainly at their centre point. This bulging effect is due to the differential pressure encountered by the holder once it is inserted in the vacuum of the microscope.. The bulging phenomenon of the windows consequently adds some extra thickness to the observation chamber, typically up to 150nm in excess per chip, to the 200nm thickness provided by the spacer. The bulging effect is at minimum however at the corners of the rectangular windows. Thus, ferritin imaging was performed by the areas close to the corners of the SixNy window. Nonetheless, the thickness of the liquid layer resides in the nanometre range, allowing for satisfactory electron penetration and sample contrast.

### Fluid dynamics simulation

Section Liquid cell preparation and assembly shows how demanding this technique can be, and it is not surprising that the first attempts to image samples in liquid state failed due the high complexity of the system. Therefore, a fluid dynamics simulation was performed by using COMSOL Multiphysics in order to understand the functioning of the liquid cell in “in-flow” conditions and obtain the best performance from the imaging process. The liquid holder was modelled in COMSOL using the actual dimensions of the device, the values of which will not be reported in this paper so as not to violate the confidentiality agreement signed between the authors and the manufacturing company. However, the simplified geometry of the holder is shown in **Fig.S1a**. The tip of the holder is flower-shaped and is connected to two in plane parallel pipes, an inlet and outlet pipe for flowing in and out of the solution respectively. The holder hosts on the inside two SiN chips separated by a variable gap, creating a microfluidic channel to enable inflow analysis and keep the sample hydrated. The bulging effect on the observation windows occurring when imaging the sample and mentioned in section Liquid cell preparation and assembly of SI has been excluded from the simulation. This is because the simulation has been performed at atmospheric pressure conditions, *i*.*e*. when the sample is flowed in the liquid holder outside the microscope. Thus, the windows are treated as flat, rigid surfaces trapping the solution on the inside. Input and output ports were set to the end of the pipes, guaranteeing good fluid circulation. At the beginning of the simulation the chamber was modelled as empty, both chips were removed, and an uncompressible Newtonian fluid was pumped in with an initial velocity of 5¬µL/min. The analysis was performed at steady state, imposing laminar flow, planar symmetry and no slip condition on the walls of the holder. The velocity of the fluid in these conditions was analyzed and the results are reported in **Fig. S1b**. This first preliminary simulation was mainly performed to validate the goodness of the model, and especially to analyze the trajectory of the flow. The first simulation confirmed that the fluid linearly went from the inlet to the outlet pipes, slowly expanding in the whole volume of the chamber. The results showed that the trajectory of the injected particles were a combination of both convection and Brownian motion experienced by nanoparticles in liquid. Then, the complexity of the system was increased by adding the two chips inside the flower-shaped tip, separated by a 200nm spacer. The inclusion of the chips significantly influenced the liquid flow inside the chamber, modifying the liquid trajectory and narrowing it down to the small microfluidic channel between the chips. Consequently, most of the fluid went into the petals of the structure, generating a turbulent flow. Furthermore, the volume between the two chips was filled with only a small portion of the incoming fluid, the motion of which was hardly moved by the effect of the pump (**Fig. S1c**). In a first approximation model, these results can be extended to the particles in solution pumped in the inflow port; mirroring the actual experimental conditions. As a consequence, most of the particles would not flow between the observation window, but instead be trapped in the flower petal reservoir. These findings strongly suggest that parameters such as sample concentration and velocity at which the sample is injected into the holder have to be carefully chosen, in order to maximize the number of the particles in the viewing window area.

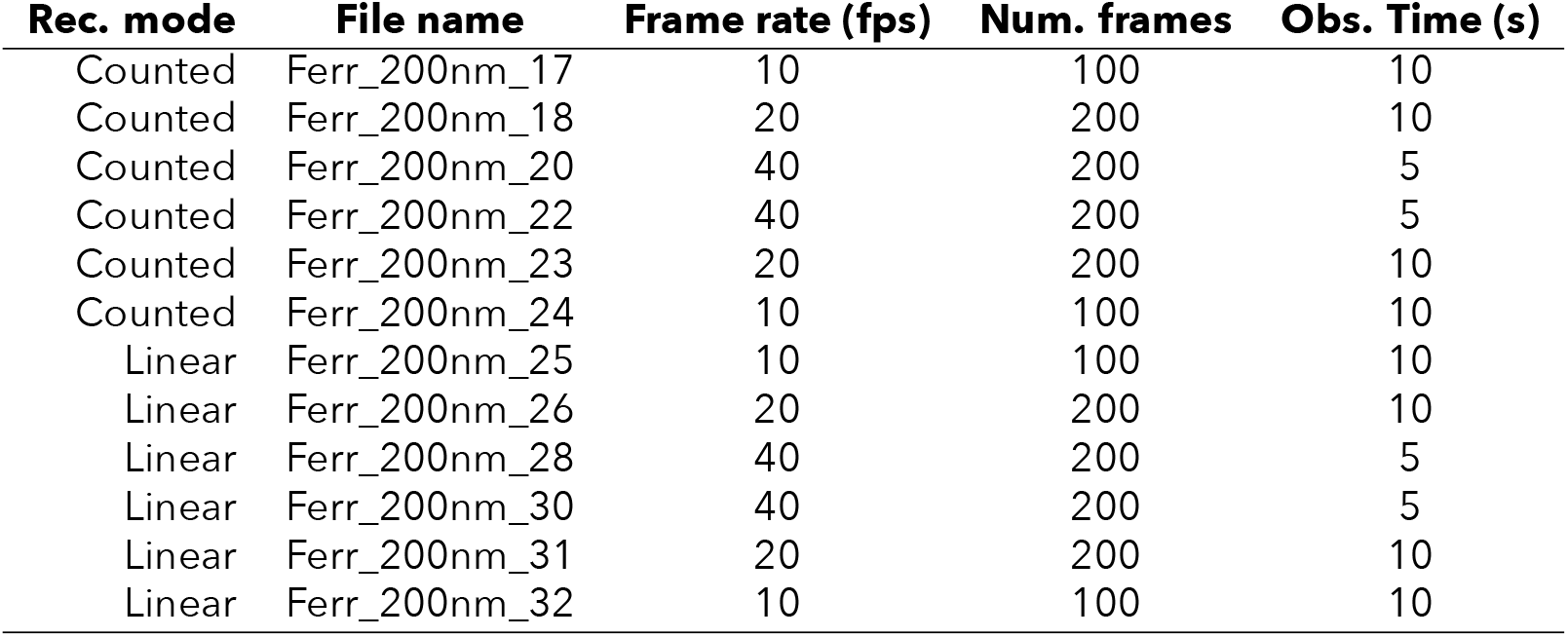

### Video analysis

The reconstruction process of the 3D density map of ferritin in solution recorded by the K2 in dose fractionation video mode has been analyzed and discussed above. However, not all the obtained videos could be used as input to the workflow.Since the sample is in a liquid state, and the particles moved with time, and the same particles could not image twice. Even with a standardised recording method, the quality of the movies could vary. In this section, 12 different videos recorded during the same session are analyzed. Table S5 contains information about the recording mode, the frame rate, the number of frames and the observation time for each video.

Figure S5, Äì Metadata extracted from the videos of ferritin in solution.

The recording mode has been widely discussed in section LP EM Imaging procedure, therefore, in this section other additional features are discussed. The file name of the videos is self-explanatory, as it contains the type of sample in solution, the size of the spacer between the chips in the holder and an identification progressive index. We processed the images, using the progressive image denoising (PID) algorithm [46]. The selection of the PID algorithm relies on its ability to produce enhanced results, without generating artefacts. However, this algorithm is very CPU demanding and time consuming; it takes about 30 minutes to process a single frame made of 1919×1855 pixels. Consequently, only a subset of three frames for every video has been selected, these frames were selected at the very beginning, in the middle and at the very end (**Fig. S6**) of each video. Running any quantitative analysis on these data may be not very accurate, due to the uncertain efficiency of this method. Yet, it is possible to notice how the counted mode generally produces better results than the linear mode and that the application of the median filter significantly increases the quality of the images.

### Performance analysis

The whole workflow was executed on a machine running Ubuntu Linux 16.04 64-bit operative system with the following hardware specifications: two central processing units (CPU) Intel Xeon Gold 5118 2.3GHz with 12 cores each, 128GB (8×16GB) 2666MHz DDR4 RAM, 512GB class 20 solid state drive (SSD) and dual SLI NVIDIA Quadro P5000 16GB as GPUs. Table S3 reports the processing time values associated to the single steps of the workflow with respect to the processed videos.

Processing time

Table S3, Äì Processing time values employed in the various steps of the workflow.

| Task | Processing time |
| --- | --- |
| MATLAB pre-processing (per frame) | 00h 00,Äô 08,Äô,Äô |
| Pre-process micrographs | 00h 04,Äô 29,Äô,Äô |
| CTF correction | 00h 02,Äô 48,Äô,Äô |
| Particles picking | 00h 03,Äô 45,Äô,Äô |
| Particles extraction | 00h 00,Äô 55,Äô,Äô |
| 3D initial map | 00h 05,Äô 02,Äô,Äô |
| 3D refinement | 02h 38,Äô 03,Äô,Äô |
| Local resolution | 00h 06,Äô 54,Äô,Äô |

**Figure S1:**
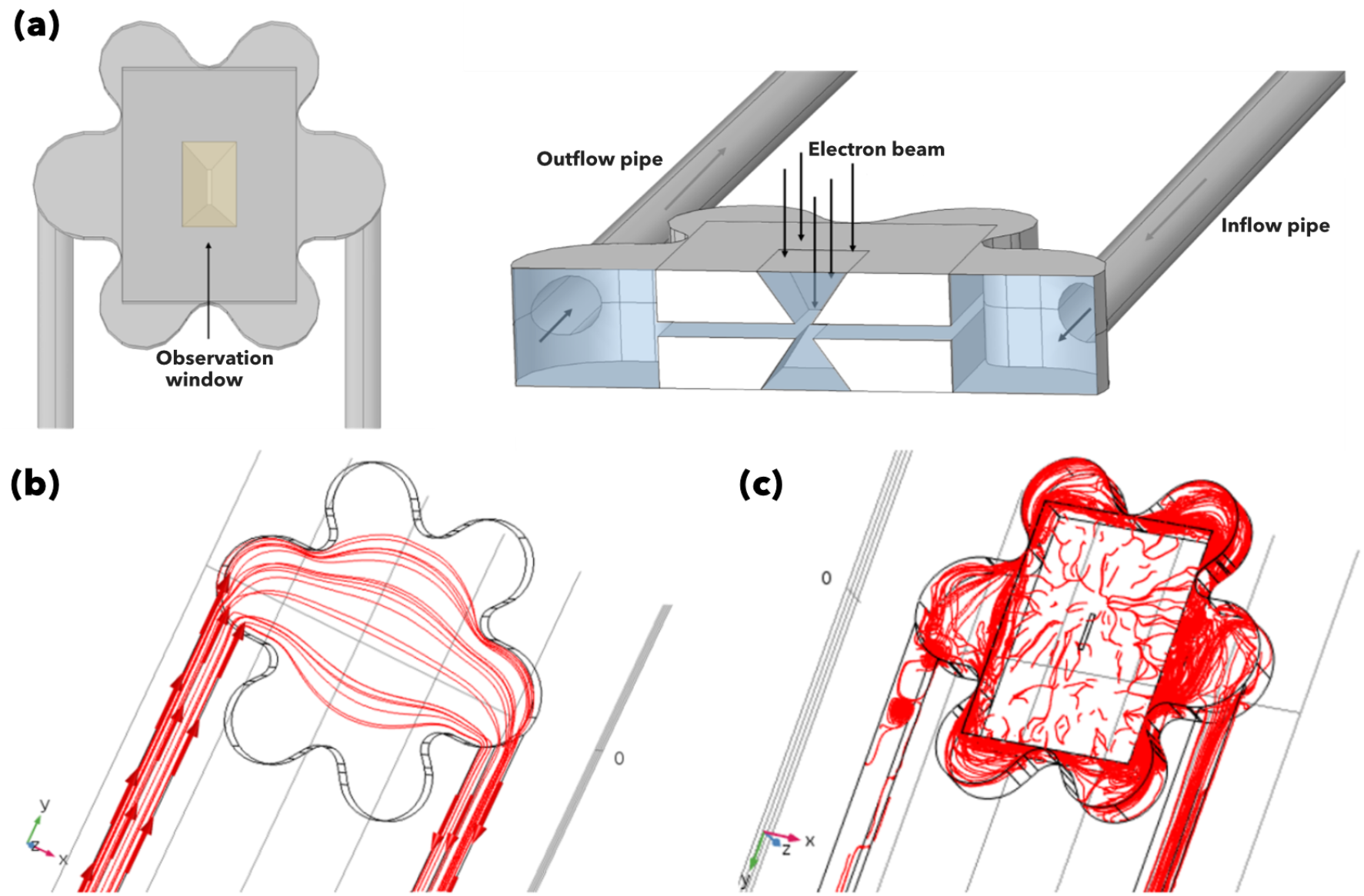
TEM holder. **(a)**Schematics of the liquid holder used for imaging liquid sample by transmission electron microscopy. **(b)** Fluid dynamics simulations of the holder, modelled without the SiN chips. A Newtonian fluid was pumped in the inflow pipe (on the left) and out from the outflow pipe (on the right). The flow expands towards the flower shape reservoir perpendicular to the liquid main trajectory while flowing through the chamber. **(c)** Fluid dynamics simulations of the holder comprising two chips inside. Both simulations run applying laminar flow, planar symmetry and no slip conditions; the analysis was then performed at steady state.

**Figure S2:**
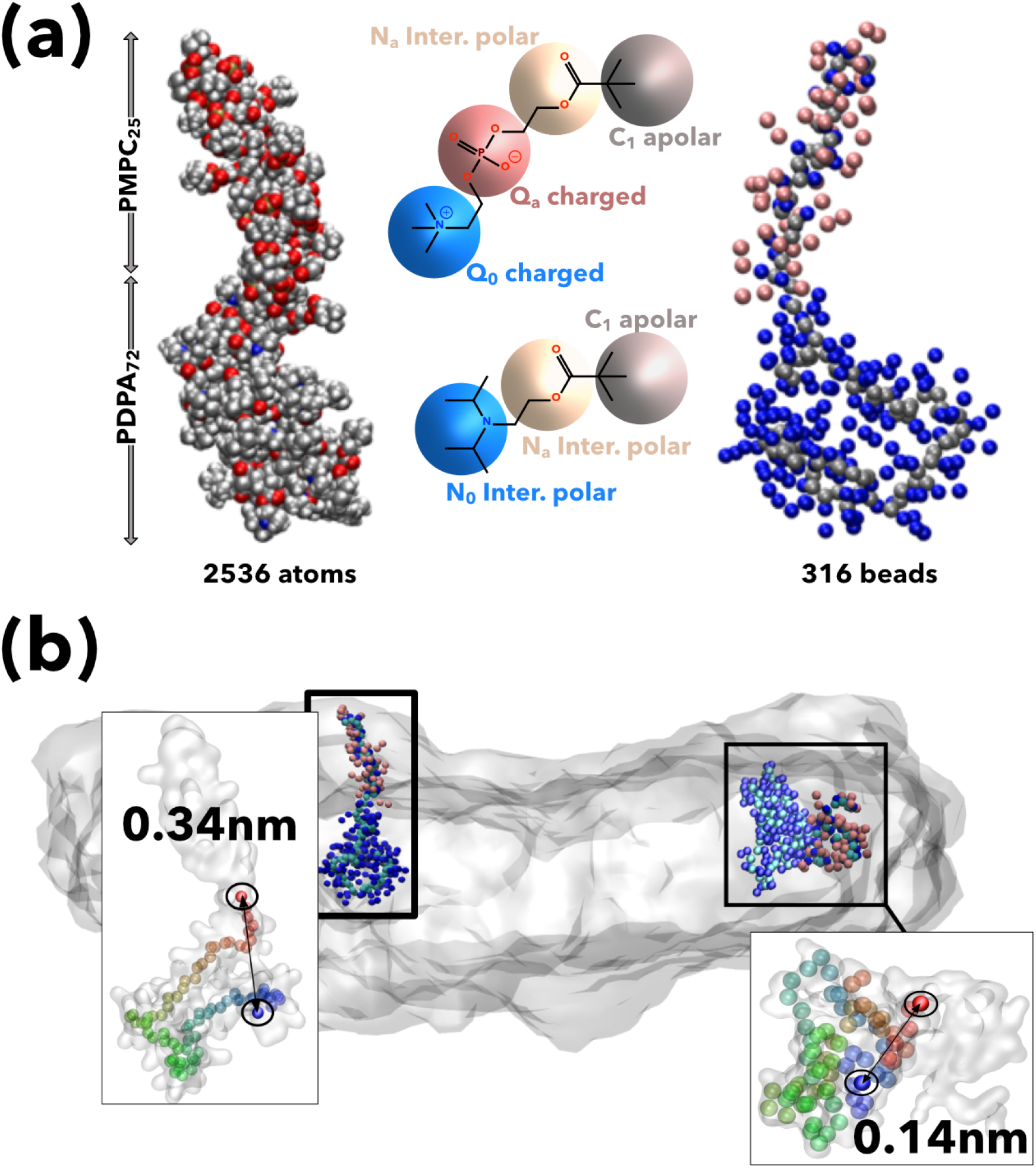
Coarse-grain model of the PMPC-PDPA micelle in water. **(a)** Coarsening of a single PMPC-PDPA chain from 2536 atoms to 316 beads. **(b)** End-to-end distance of a PMPC-PDPA chain from the core and edge areas of the the disk-like micelle.

**Figure S3:**
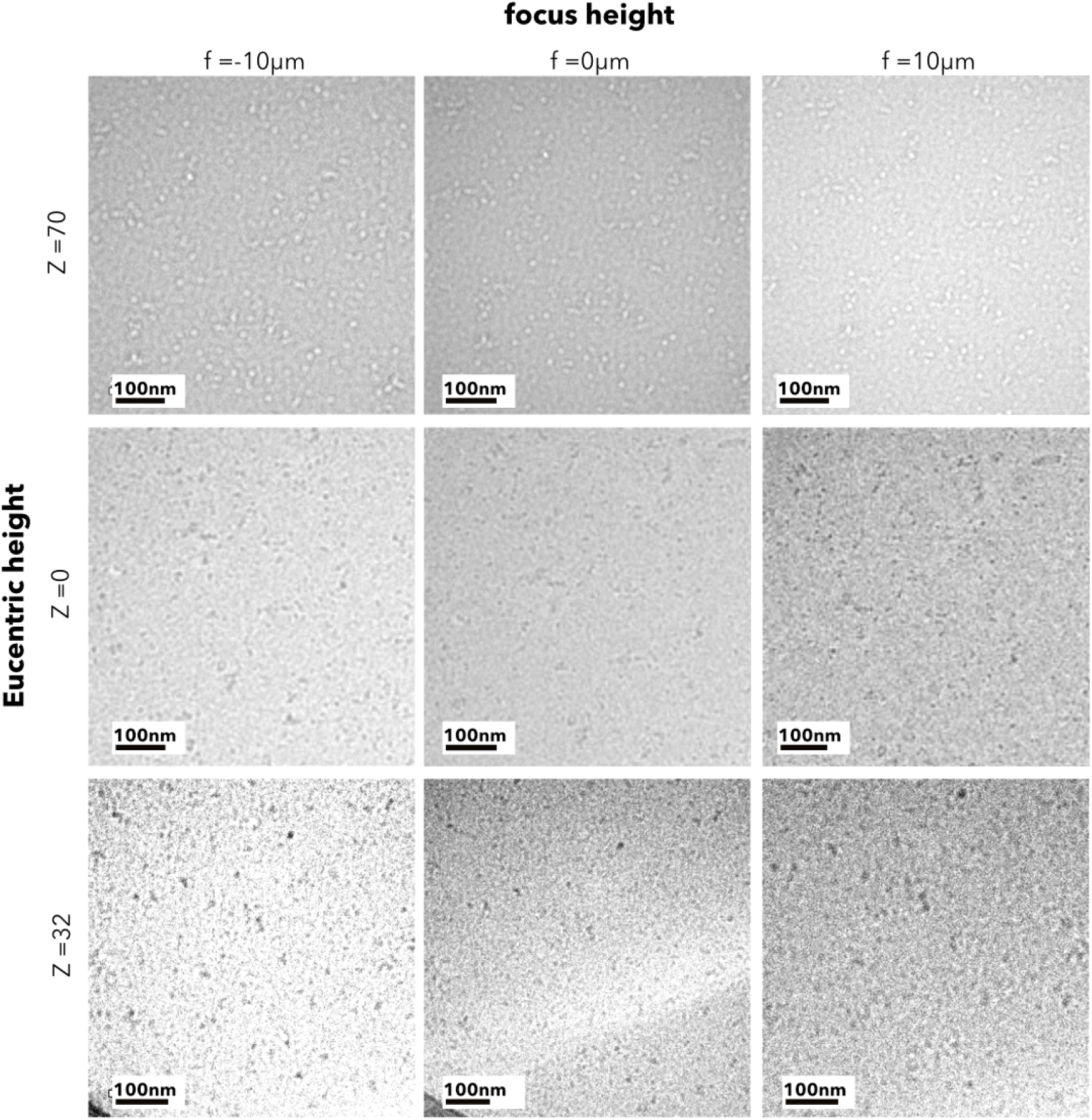
Focus and eucentric height. LPEM micrographs of ferritin protein dispersed in PBS taken at different focus and eucentric height

**Figure S4:**
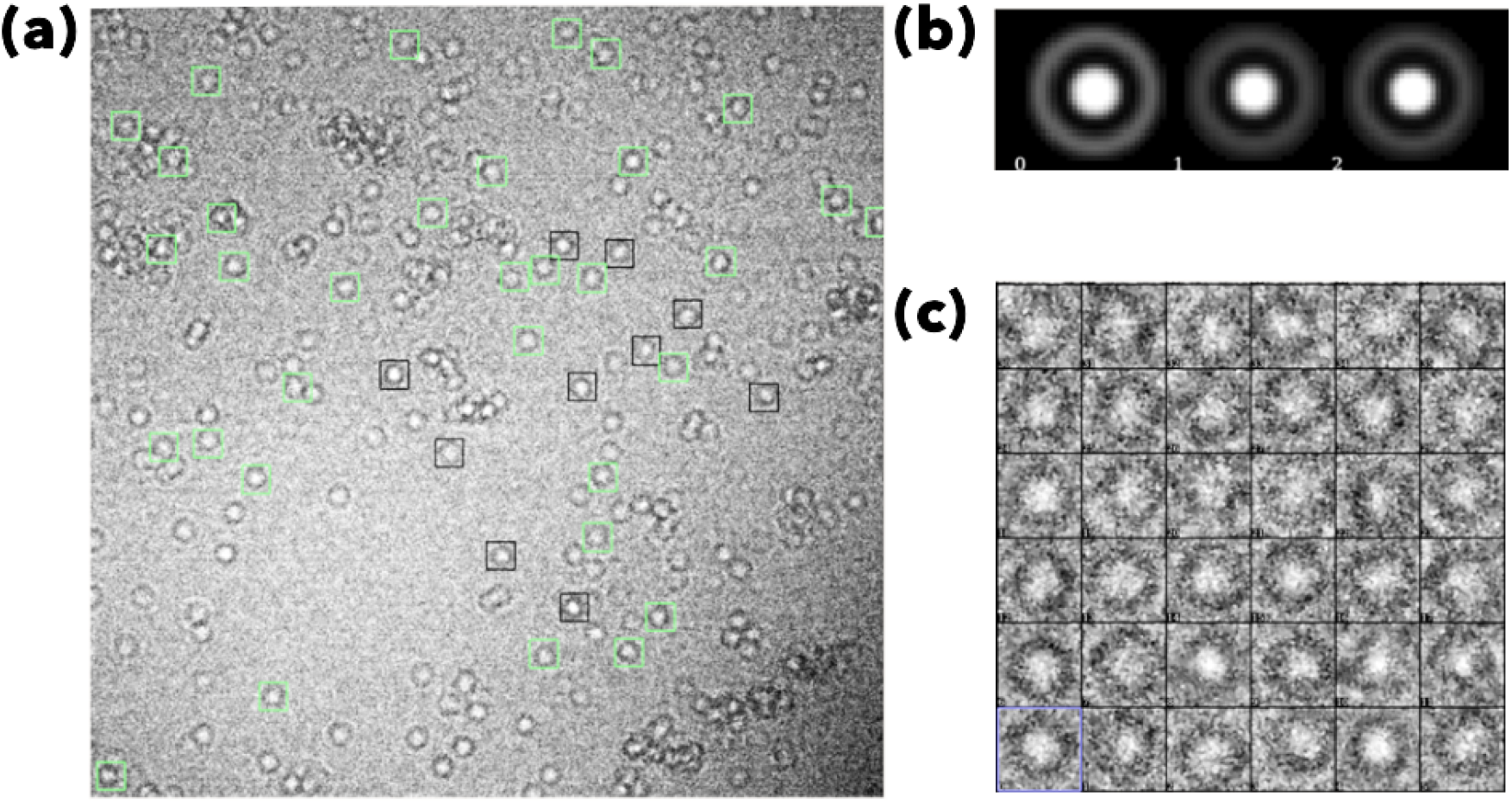
Schematic representation of the ferritin particle selection process. **(a)** The black boxes represent the ferritin particles selected by the user, which are transformed into a template. **(b)** The template is then used to recognise and identify all the occurrences in the rest of the frame (in green). **(c)** The particles automatically selected are saved into a gallery. Lastly, the so-defined particle identification scheme is applied to the rest of the frames comprising the videos.

**Figure S5:**
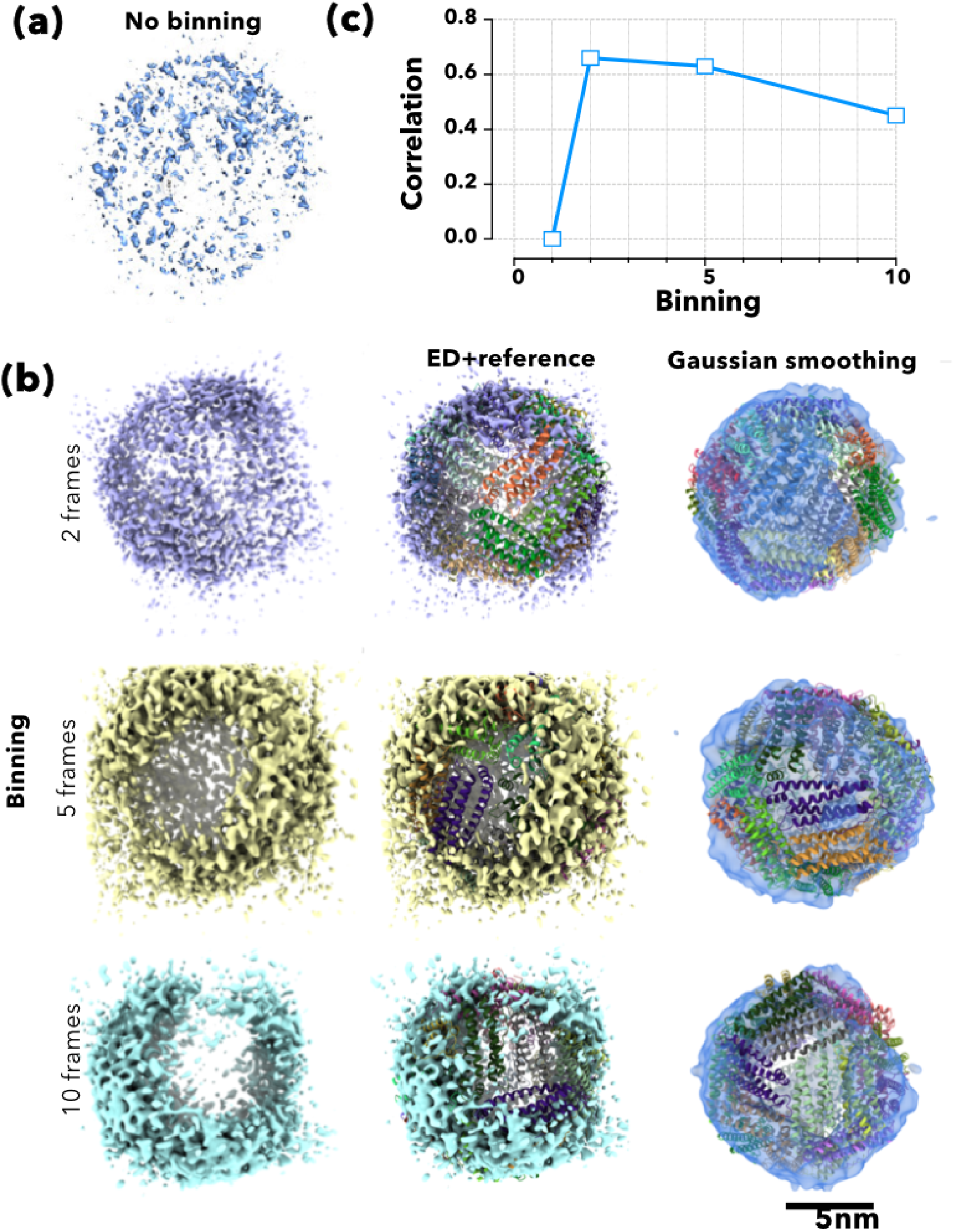
Low dose 3D rendering. 3D renderings of the ferritin imaged a ultra low dose and reconstructed using different level of binning **(a)**. Comparison between the low dose renderings showed as no processing and as Gaussian-smoothed and reference x-ray structure showed as overlays between structures **(b)** and the corresponding correlation **(c)**.

**Figure S6:**
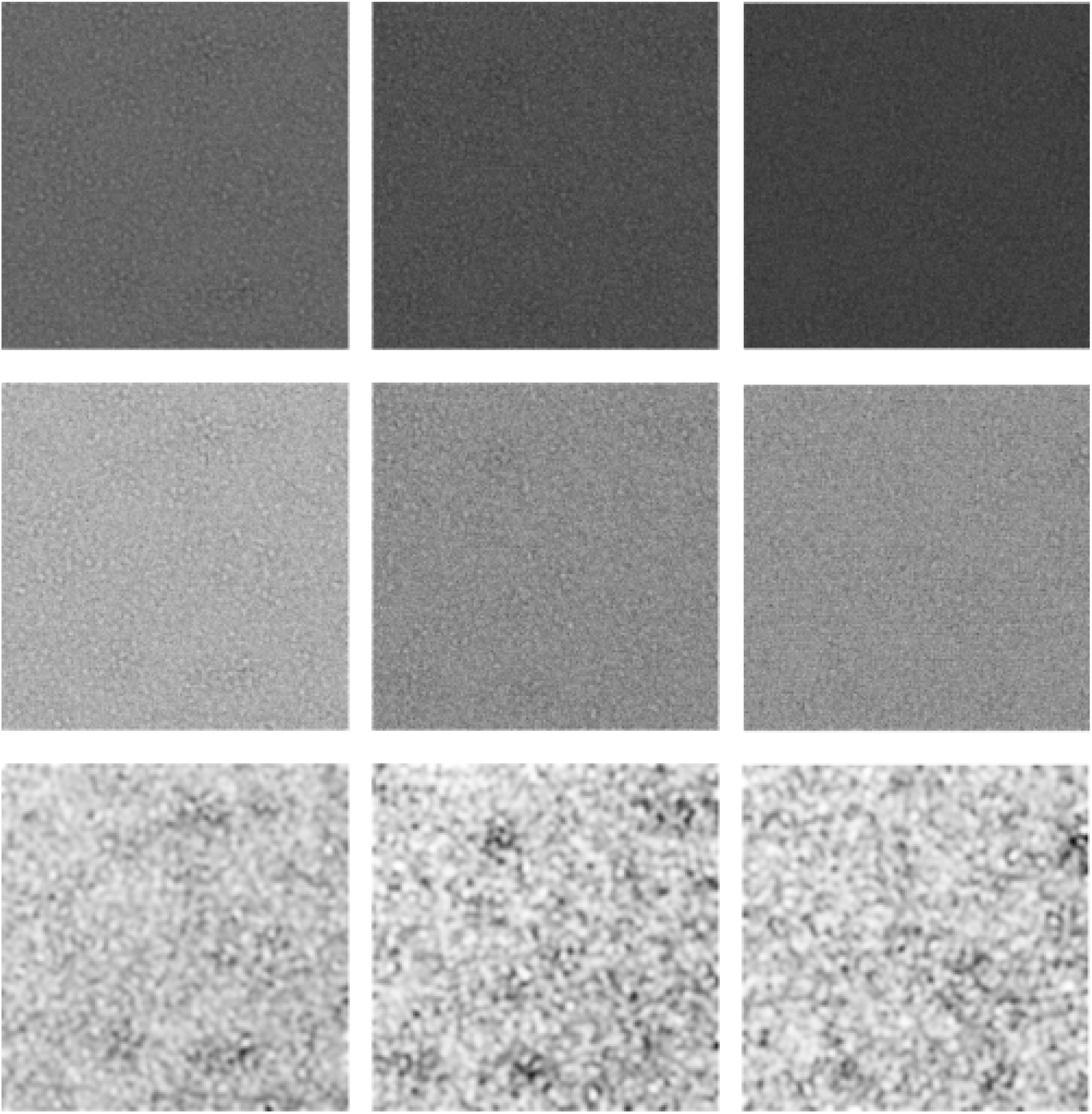
Examples of frames to show denoising . The columns of this matrix disposition represent the different frame rate values: from left to right, 10fps, 20fps and 40fps. The rows contain the different element of the analysis: from top to bottom, the original frame, the median filtered version of the original image and the reconstructed ground truth. The noiseless image at the bottom was obtained by applying the PID algorithm to the original image.

**Figure S7:**
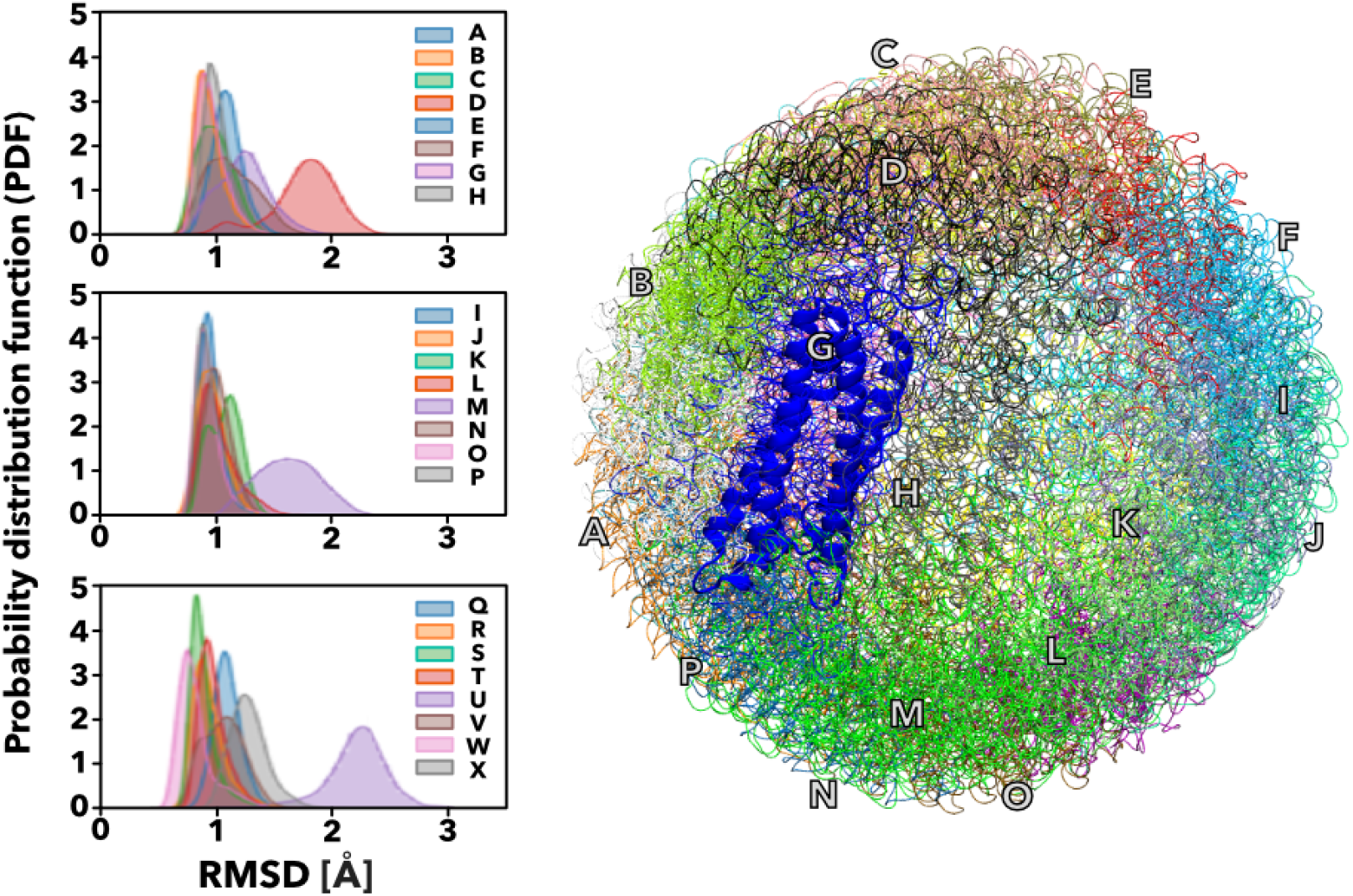
RMSD analysis. Probability distribution function (PDF) of the root mean square deviation (RMSD), along the production runs, per chain calculated from Molecular dynamic simulations of horse-spleen ferritin (PDBid 6MSX) light chain depicted as cartoon, the 24 chains are represented as letters of the alphabet.

## References

[1] AW McDowall, JJ Chang,R Freeman, J Lepault, CA Walter, andJ Dubochet. Electron microscopy of frozen hydrated sections of vitreous ice and vitrified biological samples. Journal of microscopy, 131:1–9, July 1983.

[2] K Koga, H Tanaka, and XC Zeng. First-order transition in confined water between highdensity liquid and low-density amorphous phases. Nature, 408:564–567, November 2000.

[3] Jacques Dubochet. The physics of rapid cooling and its implications for cryoimmobilization of cells. Methods in cell biology, 79:7–21, 2007.

[4] Jacques Dubochet. On the development of electron cryo-microscopy (nobel lecture). Angewandte Chemie (International ed. in English), 57:10842–10846, August 2018.

[5] Joachim Frank. Single-particle reconstruction of biological molecules-story in a sample (nobel lecture). Angewandte Chemie (International ed. in English), 57:10826–10841, August 2018.

[6] Richard Henderson. From electron crystallography to single particle cryoem (nobel lecture). Angewandte Chemie (International ed. in English), 57:10804–10825, August 2018.

[7] Xiao-chen Bai, Greg McMullan, and Sjors H W Scheres. How cryo-em is revolutionizing structural biology. Trends in biochemical sciences, 40:49–57, January 2015.

[8] Yifan Cheng, Robert M Glaeser, and Eva Nogales. How cryo-em became so hot. Cell, 171:1229–1231, November 2017.

[9] J Frank, A Verschoor, and M Boublik. Computer averaging of electron micrographs of 40s ribosomal subunits. Science (New York, N.Y.), 214:1353–1355, December 1981.

[10] Yifan Cheng. Single-particle cryo-em-how did it get here and where will it go. Science (New York, N.Y.), 361:876–880, August 2018.

[11] Richard Henderson. Realizing the potential of electron cryo-microscopy. Quarterly reviews of biophysics, 37:3–13, February 2004.

[12] Alan Merk, Alberto Bartesaghi, Soojay Banerjee, Veronica Falconieri, Prashant Rao, Mindy I Davis, Rajan Pragani, Matthew B Boxer, Lesley A Earl, Jacqueline L S Milne, and Sriram Subramaniam. Breaking cryo-em resolution barriers to facilitate drug discovery. Cell, 165:1698–1707, June 2016.

[13] Ahmed H Zewail. Four-dimensional electron microscopy. Science (New York, N.Y.), 328:187–193, April 2010.

[14] Niels de Jonge and Frances M Ross. Electron microscopy of specimens in liquid. Nature nanotechnology, 6:695–704, October 2011

[15] Aleksandar Radisic, Philippe M Vereecken, James B Hannon, Peter C Searson, and Frances M Ross. Quantifying electrochemical nucleation and growth of nanoscale clusters using real-time kinetic data. Nano letters, 6:238–242, February 2006.

[16] Zhiyuan Zeng, Wen-I Liang, Hong-Gang Liao, Huolin L Xin, Yin-Hao Chu, and Haimei Zheng. Visualization of electrode-electrolyte interfaces in lipf6/ec/dec electrolyte for lithium ion batteries via in situ tem. Nano letters, 14:1745–1750, 2014.

[17] Haimei Zheng, Rachel K Smith, Young-Wook Jun, Christian Kisielowski, Ulrich Dahmen, and A Paul Alivisatos. Observation of single colloidal platinum nanocrystal growth trajec-tories. Science (New York, N.Y.), 324:1309–1312, June 2009.

[18] MJ Williamson, RM Tromp, PM Vereecken, R Hull, and FM Ross. Dynamic microscopy of nanoscale cluster growth at the solid-liquid interface. Nature materials, 2:532–536, August 2003.

[19] Stephan Thiberge, Amotz Nechushtan, David Sprinzak, Opher Gileadi, Vered Behar, Ory Zik, Yehuda Chowers, Shulamit Michaeli, Joseph Schlessinger, and Elisha Moses. Scanning electron microscopy of cells and tissues under fully hydrated conditions. Proceedings of the National Academy of Sciences of the United States of America, 101:3346–3351, March 2004

[20] . Diana B Peckys, Gabriel M Veith, David C Joy, and Niels de Jonge. Nanoscale imaging of whole cells using a liquid enclosure and a scanning transmission electron microscope. PloS one, 4:e8214, December 2009.

[21] Jungwon Park, Hans Elmlund, Peter Ercius, Jong Min Yuk, David T Limmer, Qian Chen, Kwanpyo Kim, Sang Hoon Han, David A Weitz, A Zettl, and A Paul Alivisatos. Nanoparticle imaging. 3d structure of individual nanocrystals in solution by electron microscopy. Science (New York, N.Y.), 349:290–295, July 2015.

[22] Laura Rodríguez-Arco, Alessandro Poma, Lorena Ruiz-Pérez, Edoardo Scarpa, Kamolchanok Ngamkham, and Giuseppe Battaglia. Molecular bionics - engineering biomaterials at the molecular level using biological principles. Biomaterials, 192:26–50, February 2019.

[23] Lucas R Parent, Evangelos Bakalis, Abelardo Ramírez-Hernández, Jacquelin K Kammeyer, Chiwoo Park, Juan de Pablo, Francesco Zerbetto, Joseph P Patterson, and Nathan C Gianneschi. Directly observing micelle fusion and growth in solution by liquid-cell transmission electron microscopy. Journal of the American Chemical Society, 139:17140–17151, November 2017.

[24] Mollie A Touve, C Adrian Figg, Daniel B Wright, Chiwoo Park, Joshua Cantlon, Brent S Sumerlin, and Nathan C Gianneschi. Polymerization-induced self-assembly of micelles observed by liquid cell transmission electron microscopy. ACS central science, 4:543–547, May 2018.

[25] Alessandro Ianiro, Hanglong Wu, Mark M J van Rijt, M Paula Vena, Arthur DA Keizer, A Catarina C Esteves, Remco Tuinier, Heiner Friedrich, Nico A J M Sommerdijk, and Joseph P Patterson. Liquid-liquid phase separation during amphiphilic self-assembly. Nature chemistry, 11:320–328, April 2019.

[26] Xuewen Fu, Bin Chen, Jau Tang, Mohammed Th Hassan, and Ahmed H Zewail. Imaging rotational dynamics of nanoparticles in liquid by 4d electron microscopy. Science (New York, N.Y.), 355:494–498, February 2017.

[27] A Cameron Varano, Amina Rahimi, Madeline J Dukes, Steven Poelzing, Sarah M McDonald, and Deborah F Kelly. Visualizing virus particle mobility in liquid at the nanoscale. Chemical communications (Cambridge, England), 51:16176–16179, November 2015.

[28] E. C. Theil. Ferritin: structure, gene regulation, and cellular function in animals, plants, and microorganisms. Annual review of biochemistry, 56:289–315, 1987.

[29] Canhui Wang, Qiao Qiao, Tolou Shokuhfar, and Robert F Klie. High-resolution electron microscopy and spectroscopy of ferritin in biocompatible graphene liquid cells and graphene sandwiches. Advanced materials (Deerfield Beach, Fla.), 26:3410–3414, June 2014.

[30] G McMullan,S Chen, R Henderson, and AR Faruqi. Detective quantum efficiency of electron area detectors in electron microscopy. Ultramicroscopy, 109(9):1126–1143, 2009.

[31] Xueming Li, Paul Mooney, Shawn Zheng, Christopher R Booth, Michael B Braunfeld, Sander Gubbens, David A Agard, and Yifan Cheng. Electron counting and beam-induced motion correction enable near-atomic-resolution single-particle cryo-em. Nature methods, 10(6):584, 2013.

[32] Claude Knaus and Matthias Zwicker. Progressive image denoising. 23:3114–3125, 2014.

[33] Jacques Cazaux. Correlations between ionization radiation damage and charging effects in transmission electron microscopy. Ultramicroscopy, 60:411–425, 1995.

[34] Manikandan Karuppasamy, Fatemeh Karimi Nejadasl, Milos Vulovic, Abraham J. Koster, and Raimond B. G. Ravelli. Radiation damage in single-particle cryo-electron microscopy: effects of dose and dose rate. Journal of synchrotron radiation, 18:398–412, May 2011.

[35] Claudia Contini, Russell Pearson, Linge Wang, Lea Messager, Jens Gaitzsch, Loris Rizzello, Lorena Ruiz-Perez, and Giuseppe Battaglia. Bottom-up evolution of vesicles from disks to high-genus polymersomes. iScience, 7:132–144, September 2018.

[36] Pierre-Gilles de Gennes. Scaling Concepts in Polymer Physics. Cornell University Press, 1979.

[37] Giuseppe Battaglia and Anthony J Ryan. Bilayers and interdigitation in block copolymer vesicles. Journal of the American Chemical Society, 127:8757–8764, June 2005.

[38] S Jonić, COS Sorzano, and N Boisset. Comparison of single-particle analysis and electron tomography approaches: an overview. Journal of microscopy, 232:562–579, December 2008.

[39] JM de la Rosa-Trevín, A Quintana, L Del Cano, A Zaldívar, I Foche, J Gutiérrez, J Gómez-Blanco, J Burguet-Castell, J Cuenca-Alba,V Abrishami, J Vargas, J Otón, G Sharov, JL Vilas, J Navas,P Conesa, M Kazemi, R Marabini, COS Sorzano, and JM Carazo. Scipion: A software framework toward integration, reproducibility and validation in 3d electron microscopy. Journal of structural biology, 195:93–99, July 2016.

[40] Alp Kucukelbir, Fred J Sigworth, and Hemant D Tagare. Quantifying the local resolution of cryo-em density maps. Nature methods, 11:63–65, January 2014.

[41] Christopher J. Russo and Lori A. Passmore. Electron microscopy: Ultrastable gold substrates for electron cryomicroscopy. Science (New York, N.Y.), 346:1377–1380, December 2014.

[42] Thomas D. Goddard, Conrad C. Huang, Elaine C. Meng, Eric F. Pettersen, Gregory S. Couch, John H. Morris, and Thomas E. Ferrin. Ucsf chimerax: Meeting modern challenges in visualization and analysis. Protein science: a publication of the Protein Society, 27:14–25, January 2018.

[43] Thomas D. Goddard, Conrad C. Huang, and Thomas E. Ferrin. Visualizing density maps with ucsf chimera. Journal of structural biology, 157:281–287, January 2007.

[44] Richard Henderson and Greg McMullan. Problems in obtaining perfect images by single-particle electron cryomicroscopy of biological structures in amorphous ice. Microscopy (Oxford, England), 62:43–50, February 2013.

[45] Christopher J Russo and Lori A Passmore. Electron microscopy: Ultrastable gold substrates for electron cryomicroscopy. Science (New York, N.Y.), 346:1377–1380, December 2014.

[46] Rafael C. Gonzalez and Richard E. Wood. Digital Image Processing. Pearson, 1192, 4th edition, 2008.

